# New distributional opportunities with niche innovation in Eurasian snowfinches

**DOI:** 10.1101/2021.04.06.438738

**Authors:** Marlon E Cobos, Yalin Cheng, Gang Song, Fumin Lei, A. Townsend Peterson

## Abstract

This study explores the evolutionary history of ecological niche characters in the Eurasian snowfinch lineage. Specifically, we use new analytical approaches to reconstruct ecological niche evolution, taking uncertainty in knowledge of the ecological niche limits into account. We found an overall pattern of niche conservatism in relation to both temperature and precipitation characteristics of niches, but for one dramatic niche evolution event, in *Montifringilla nivalis*. Interestingly, this species is also that which has by far the broadest geographic distribution among snowfinches. We conclude that an evolutionary change in niche characteristics perhaps within *M. nivalis* (i.e., present in some and not all of its populations) made possible the broad, westward range expansion of that species, thus changing the distributional potential of the snowfinch lineage dramatically.

## Introduction

The field of distributional ecology is both new and old—old in the sense that biologists have been intensely interested in geographic distributions of species since pre-Darwinian times, but new in the sense that a new suite of tools and frameworks has brought new rigor and insight (Johnson et al., 2019; Randin et al., 2020; Zurell and Engler, 2019). A question of particular interest has been the frequency with which ecological niches change over phylogeny—the idea of phylogenetic niche conservatism has enabled the tools termed ecological niche modelling (Peterson et al., 2011), and inspired speculation about implications for speciation rates (Wiens and Graham, 2005). Although studies have attempted to address niche conservatism over phylogeny for the past two decades (Graham et al., 2004; Kozak and Wiens, 2006; Peterson, 2009; Peterson et al., 1999), methodologies have not always been appropriate and rigorous (Owens et al., 2020; Saupe et al., 2018; Warren et al., 2008), which has led to a general overestimation of frequencies of ecological niche innovation (Peterson, 2011; Saupe et al., 2018).

A set of novel tools has now clarified the sources of rampant bias in (over-)estimating evolutionary rates in ecological niches over phylogenies (Owens et al., 2020), permitting rigorous identification of niche innovation events. Specifically, these methods separate tolerance spectra of species into distinct bins, and score each bin as to present, absent, or uncertain, the latter category referring to situations in which no environments are available within areas accessible to species to permit conclusions of absence (Owens et al., 2020). The result is a more conservative, but apparently much less biased, view of ecological niche innovation over phylogeny in many groups (Ribeiro et al., 2016).

Perhaps most interesting is that these methods now allow identification of new distributional possibilities that open with each niche evolution event, which we explore in this paper. The snowfinches are a lineage of Old World sparrows that range across Eurasia, mostly in remote and high-montane regions, which has seen detailed phylogenetic (e.g., Qu et al., 2006; Qu and Lei, 2009; Yang et al., 2006) and genomic (Qu et al., in review) study. Here, we apply novel phylogenetic-reconstruction tools to the ecological niches of snowfinches, and explore the distributional implications of the niche changes that have indeed occurred in the history of this group. This first application and exploration of such questions in a relatively simple lineage presages a broader and more comprehensive understanding of the changing distributional potential of lineages through time.

## Materials and Methods

### Occurrence data

Occurrence data for the snowfinch clade were obtained from eBird (https://ebird.org/; Sullivan et al., 2009), via searches for *Montifringilla adamsi*, *M. nivalis*, *M. henrici*, *M. taczanowskii*, *M. blanfordi*, *M. ruficollis*, *M. davidiana*, and *M. theresae*, although these taxa are often treated as belonging to three genera (*Montifringilla*, *Onychostruthus*, and *Pyrgilauda*; Qu et al., 2006). We filtered the initial eBird query to remove long-distance, broad-spatial-footprint reports for which geographic referencing would be imprecise by removing traveling counts for which traveling distance was >10 km, and other protocols for which no traveling distance was given. The initial count of occurrence data available across all of the species was 3266; however, after removing duplicate records and keeping only one from each set of records separated by <10 km, we had only 1353 records remaining. Sample sizes per species at this point ranged 41–805 (Fig. 1). For *Pyrgilauda theresae*, only two records were available; because these two sites were closely similar in environmental characteristics, and in light of the apparent close relationship of this species with *P. blanfordi* (with which it has been considered conspecific; BirdLife International, 2020) but complete lack of high-quality tissue samples, we removed this species from the analysis.

**Fig. 1.**
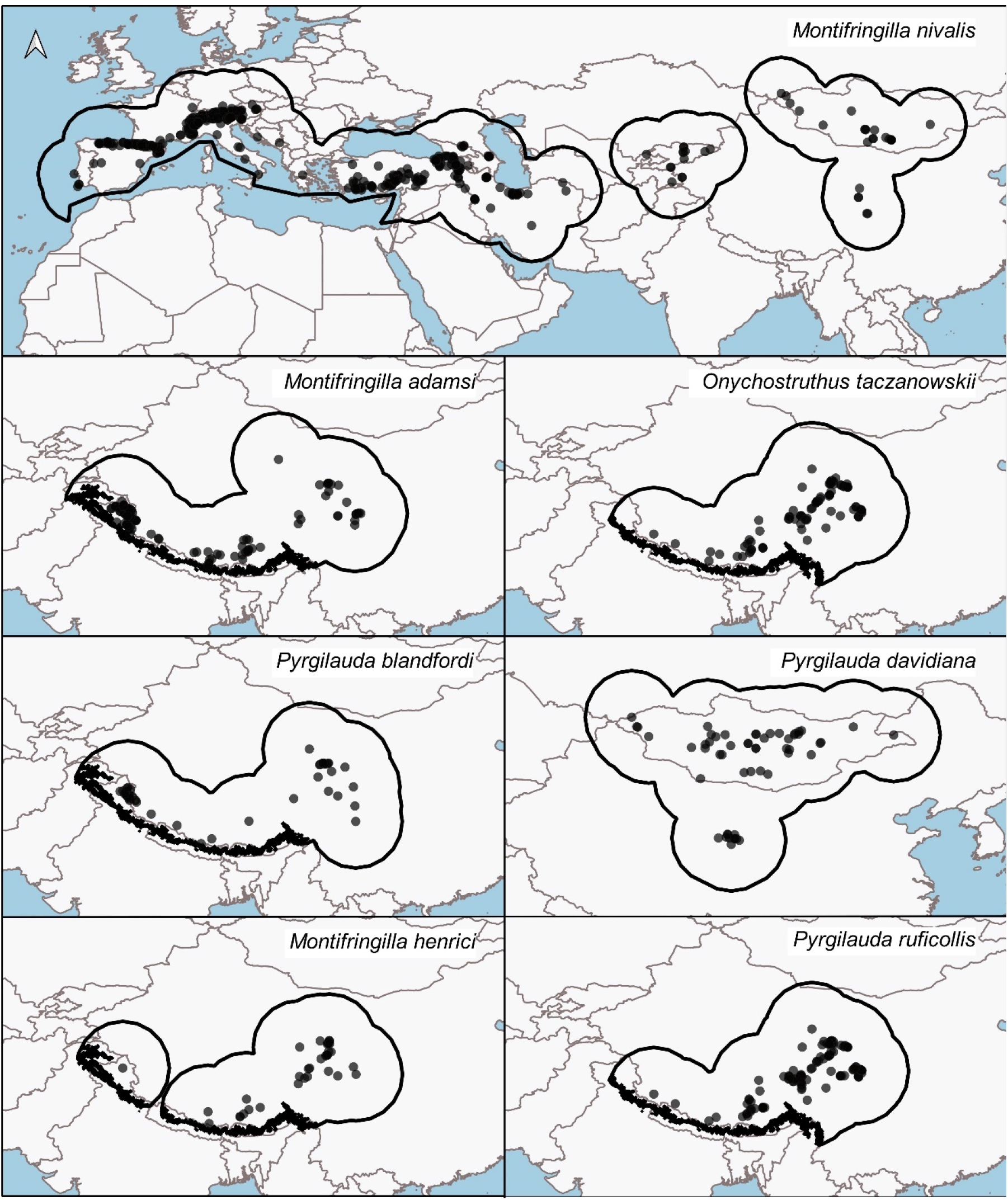
Occurrence records and hypotheses of areas that have been accessible to each species over relevant time periods (M) for the seven species of snowfinches of interest.

### Climate data

Data summarizing variation in climatic dimensions across Eurasia were obtained from the Chelsa climate database (http://chelsa-climate.org/), which summarize global climates over the period 1979–2013 (Karger et al., 2017). Data layers had a spatial resolution of 30”, or approximately 1 km at the Equator. We excluded from analysis four variables that combine temperature and precipitation information (bio 8, bio 9, bio 18, bio 19), to avoid problems deriving from odd spatial artefacts (Escobar et al., 2014). Variables used can be obtained using the code provided in Supplementary Data, file name: Commented_script.pdf

### Phylogenetic hypotheses

We explored initially three distinct phylogenetic hypotheses: (1) a coalescent tree constructed by ASTRALⅡ (Mirarab and Warnow, 2015) based on 2000 trees randomly extracted from birdtree (http://birdtree.org/) that contains seven species, including all snowfinch species except *M. henrici*, (2) a Bayesian consensus tree generated using BEAST2 (Bouckaert et al., 2014) based on 13 mtDNA of all snowfinch species except *P. theresae*, and (3) a coalescent tree based on maximum likelihood analyses of 13 mtDNA constructed with RAxML (Stamatakis, 2014) using ASTRALII (Supplementary Data, Fig. S1 has the summary). One species (*P. theresae*) was excluded from analysis owing to the small number of occurrence records (*n* = 2), which reduced the accuracy with which niche limits could be detected; what is more, the best phylogenetic hypothesis available to us did not include this taxon. The consensus phylogenetic tree for tree 2 (see above) was obtained from a total of 135,003 trees in the *a posteriori* distribution, among which only a single tree had a different topology; all remaining trees presented the same topology and varied only in branch length (Supplementary Data, Fig. S2). Occurrence data and phylogenetic trees are available in Supplementary Data, file name: Initial_data.zip.

### Data Analysis

We focused our data analysis on the challenge of reconstructing the evolutionary history of fundamental niches in the snowfinch radiation. Ecological niches of species were characterized in single environmental dimensions at a time—the methodology was designed and coded this way for simplicity, and because numbers of occurrence data are insufficient for multivariate analyses for many or most species. All of our analyses follow methods presented by Owens et al. (2020), following insights into the importance of considering uncertainty in such reconstructions (Saupe et al., 2018)—Owens et al. (2020) offered detailed justifications of methodological decisions (summarized below), and presented a set of tools that supports all of the analyses conducted in this study.

### Accessible area hypotheses

In light of the relatively good abilities of snowfinch species as regards movement, and exploring and detecting areas presenting suitable conditions (Brambilla et al., 2019), we used a relatively broad area for defining species-specific accessible areas (termed **M**, see Soberón and Peterson, 2005), which in turn are the areas across which niche models should be calibrated (Barve et al., 2011). We defined accessible areas for each species as the area within 5° around the occurrence records of each species. Accessible areas of species in the Himalaya region were further limited to only areas above 2500 m elevation (Fig. 1; Supplementary Data, Table S1). All of these processes were executed in QGIS 3.10.0 (QGIS Development Team, 2019). Supplementary Data (file name: Initial_data.zip) contains shapefiles of the **M** hypotheses used.

We inspected the occurrence data carefully for errors and inconsistencies. In particular, we checked the occurrences for consistencies with known ranges of each species (BirdLife International, 2020), and examined environmental characteristics of each occurrence point to detect potential outliers (Supplementary Data, file names: Histograms_joined.pdf and Ranges_joined.xlsx). We then inspected visualizations of the distributions of points with respect to each of the environmental dimensions, in which we focused on the extreme 5% of variable values inside **M** (2.5% on each side). The environmental range within **M**, which is crucial to establishing conditions under which a species is absent, was constrained to exclude rare values across **M** that either the species has not been able to explore or are so scarce that the species may not have been detected there.

### Ecological niche characters

We developed tables of character values for all species for each environmental dimension, summarizing ranges of variable values occupied by each species and manifested across each species’ **M** (Supplementary Data, file name: Character_tables_joined.xlsx). This summary of variable values in **M** and for occurrences allowed us to recognize values of the variable for which the species is present (value = 1), absent (value = 0), or its presence is uncertain (value = ?). Uncertainty was assigned in cases in which the occupied range abutted the environmental limit of the **M** area, such that at what value the species’ niche limit is manifested cannot be established (see Owens et al. 2020 for details). To permit detailed analysis and explicit consideration of uncertainty in our reconstructions, environmental ranges were divided into equal-width bins, each of which was subjected to phylogenetic reconstruction independently (see below). These methodological steps were taken using the package nichevol 0.1.19 (Cobos et al., 2020) in R 3.6.1 (R Core Team, 2019).

### Ancestral reconstruction of niches

Ancestral niche reconstructions were performed using the three phylogenetic hypotheses available for these taxa and the tables of characters described above. We used maximum parsimony (MP) and maximum likelihood (ML) methods to perform reconstructions for all 15 variables, resulting in 30 processes per phylogenetic tree. We used the nichevol package, which internally uses the function “asr_max_parsimony” from castor (Louca, 2020) and the function “ace” from ape (Paradis and Schliep, 2018), to perform MP and ML reconstructions, respectively. MP reconstructions were done with “all equal” transition costs, whereas ML reconstructions were done using the “equal rates” model.

We sampled 1 out of every 100 trees from the *a posteriori* distribution of trees (for tree 2 only), and developed MP and ML reconstructions on each of those 1350 trees to explore variability in our results. Reconstructed niche characters were categorized as 0 = absence, 0.5 = unknown, and 1 = present. To summarize variability, we averaged the 1350 outcomes for each current and ancestral species in each environmental dimension. Under this scheme, highly variable results will end in values in between the three recategorized states; whatever the case, values closer to 0.5 reflect uncertain or unknown states, whereas values close to 0 or 1 are much more certain. These analyses were performed using nichevol and base functions of R.

### Characterization of niche evolution

To detect environmental values that represent niche evolution between ancestors and descendant species, we compared reconstructed niches of ancestor species with those of corresponding descendant species to identify episodes of niche expansion, retraction, or stasis. This comparison was achieved as follows: (1) expansion is manifested as absence in ancestor and presence in descendant for a given “bin” of environmental values; (2) retraction is manifested as presence in ancestor and absence in descendant; and (3) stasis is the conclusion when states are equivalent in ancestor and descendant, and in any comparison in which no evidence indicates difference because the character of either ancestor or descendant is unknown (see details in Owens et al., 2020).

To identify geographic areas inside species’ **M** areas that are related to niche changes detected, we queried the **M** areas for environmental variable values that correspond to either niche expansion or retraction—given the results obtained. We developed such geographic assessments for the four taxa (*M. nivalis, M. henrice, P. davidiana,* and *P. blanfordi*) for which niche evolution was detected, using results obtained for tree 2 and MP reconstructions. We performed these tasks using the R packages raster 3.0.7 (Hijmans, 2019), rgdal 1.4.8 (Bivand et al., 2019), and nichevol. Maps were developed in QGIS. Code for performing all analysis presented is included in Supplementary Data, file name: Commented_script.pdf.

## Results

The snowfinch lineage is broadly distributed across Eurasia, with highest species richness on and around the Qinghai-Tibet Plateau. Of the seven species analyzed, five species show a quite-similar distributional pattern: that is, *Montifringilla adamsi*, *M. henrici*, *Onychostruthus taczanowskii*, *Pyrgilauda blandfordi*, and *P. ruficollis*, all have geographic distributions along the Himalayas, and extending northward onto the Qinghai-Tibet Plateau (Fig. 1). One exception is *P. davidiana*, which is distributed more to the north, across the Gobi Desert region of Mongolia, with a southernmost outlier population in central China. The most dramatic exception, however, is that of *M. nivalis*, which has a wide distribution mirroring that of *P. davidiana* in the east, but that then extends into the mountains of Central Asia, through the Levant, and across southern Europe.

We developed ancestral state reconstructions for 15 bioclimatic variables (i.e., all save for four that show odd spatial artefacts), but patterns in each variable were largely parallel, at least among temperature variables and precipitation variables separately. Patterns of niche evolution across distinct phylogenetic trees were also similar (especially for temperature), as a consequence, we present only our reconstructions for annual mean temperature and annual precipitation, and all of the remaining reconstructions are available in Supplementary Data, file name: Reconstruction_figures.zip. MP reconstructions for temperature (Fig. 2) were mostly highly conserved, such that only in *M. nivalis* were changes of more than a single bin of environmental values noted. In precipitation, our ancestral reconstructions (Fig. 3) showed a broad ancestral niche, which then manifested retractions in *M. henrici*, *O. taczanowskii*, *P. ruficollis*, and *P. davidiana*. ML reconstructions for the 15 bioclimatic variables were consistent and parallel with the MP reconstructions, but ancestral states less well defined, such that clear conclusions were difficult. The ML reconstructions are shown in Supplementary Data, file name: Reconstruction_figures.zip.

**Fig. 2.**
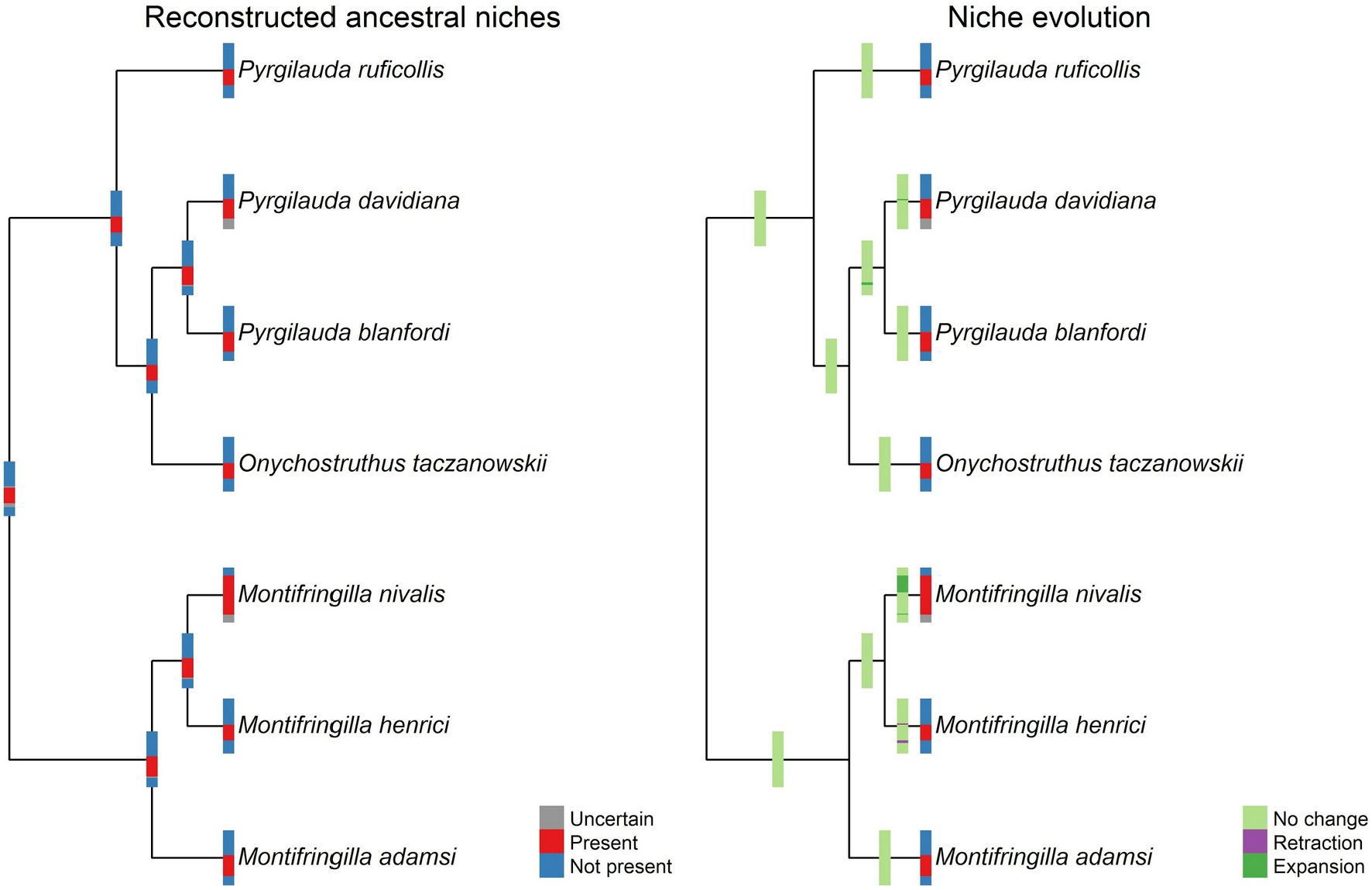
Results of maximum parsimony reconstructions of ecological niche evolution for seven species of snowfinches using annual mean temperature. Temperature is shown in a spectrum from low temperature at the bottom of the bar to high temperature for the upper part of the bar. To obtain better visualizations, the length of branches was not considered in plots (but see Supplementary Data, Fig. S1).

**Fig. 3.**
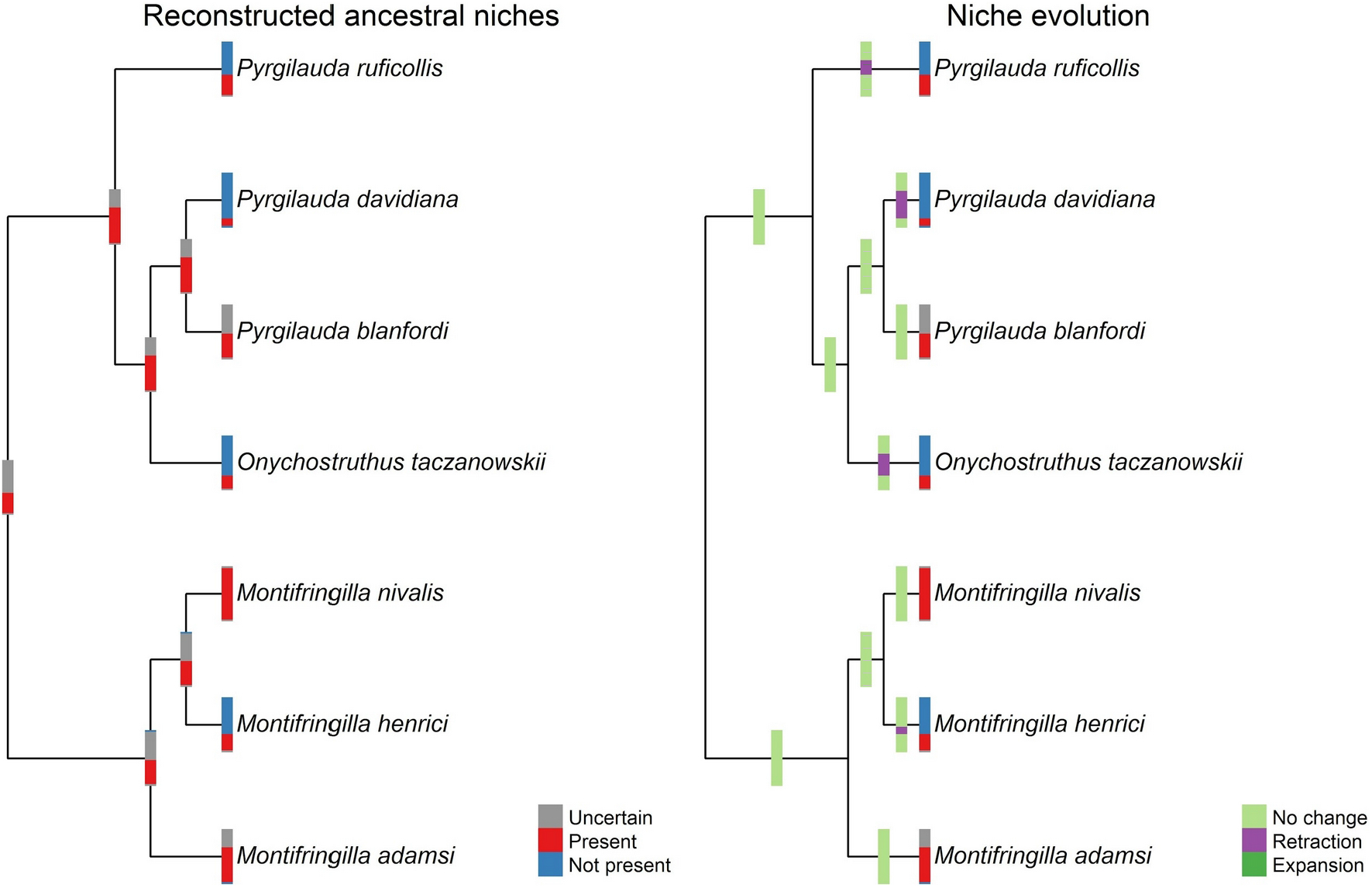
Results of maximum parsimony reconstructions of ecological niche evolution for the seven species of snowfinches using annual precipitation. Precipitation is shown in a spectrum from low precipitation at the bottom of the bar to high precipitation for the upper part of the bar. To obtain better visualizations, the length of branches was not considered in plots (but see Supplementary Data, Fig. S1).

Checking how much our ancestral state reconstructions varied among the 1350 trees sampled from the *a posteriori* distribution, we observed low variability. Indeed, for MP reconstructions, we observed no variability whatsoever in reconstructions based on different trees (which again, differed only in minor ways in terms of branch lengths). For ML reconstructions, results varied little, but no change was >0.08 along the spectrum from 0 to 1 in unsuitability to suitability (values of 0.5 representing uncertainty; Supplementary Data, file name: Variability_in_reconstructions_joined.xlsx).

Areas within the **M** for each species were queried to detect areas presenting conditions that are involved in recent niche evolution events (i.e., expansions or retractions). The most dramatic such events concerned *M. nivalis* (Fig. 4). In which the niche expansion event occurred since the split of *M. nivalis* from *M. henrici*, dated perhaps at 2.6 mya (Gebauer et al., 2006; although based on a weak molecular clock approach), such that many sectors of the western portion of the range of the species are under conditions that represent newly derived parts of the species’ ecological niche. Temperature changes in the niches of *M. henrici* and *P. davidiana* had much more subtle geographic implications (Supplementary Data, Fig. S3).

**Fig. 4.**
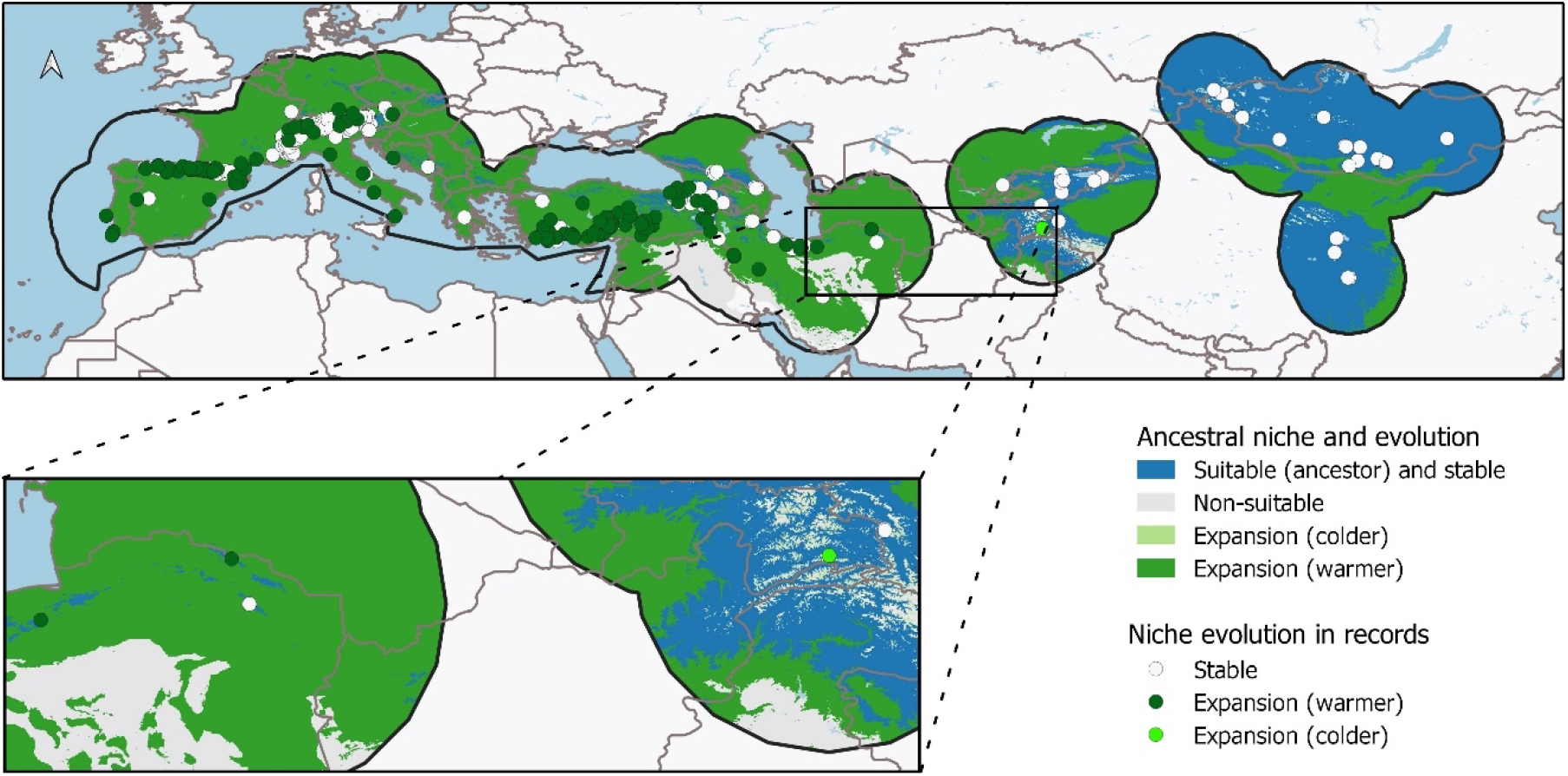
Regions inside the accessible area (**M**) of *Montifringilla nivalis* and records of this species that present conditions detected as an expansion of niche compared to its ancestor. Variable represented is the mean annual temperature. Expansion in temperature was towards higher and lower values of this variable (see Fig. 2).

The changes that we observed in niche with respect to precipitation were chiefly in the form of niche retractions (Fig. 3). These changes affected only the peripheral parts of the **M** areas for each species, as can be observed in Supplementary Data, Fig. S3. That is, in each case, areas presenting conditions from which the niche has recently retracted were along the more remote parts of the **M** area of the species.

## Discussion

The concept of the Hutchinsonian Duality (Colwell and Rangel, 2009) becomes quite significant in pondering the results that we obtained in this study. That is, species exist in two linked spaces—geographic and environmental, and they must conserve a nonzero range potential in each of those spaces to avoid extinction (Peterson, 2009). This point is partly self-evident—that is, if a species’ geographic range area approaches nil, or if conditions across that range all become unsuitable, the species will go extinct. However, the explorations presented in this study, in which we detect niche evolution *via* rigorous phylogenetic analyses that explicitly take into account uncertainty, and then examine the geographic implications, point to some subtleties. For example, the niche expansion observed in temperature tolerance of *M. nivalis* (Fig. 2) made possible massive geographic expansions in the geographic potential of the lineage (Fig. 4), yet the even larger changes in the form of niche retractions in precipitation values detected for other species, covered much smaller areas (Fig. S3).

Although analyses of ecological niche evolution in phylogenetic contexts have been developed now for more than 20 years (Graham et al., 2004; McCormack et al., 2010; Rice et al., 2003), they were rather universally compromised by lack of ability to take into account accessible areas for each species and associated uncertainty regarding suitability limits (Saupe et al., 2018). With the development of methods that are designed to take uncertainty into account explicitly (Owens et al., 2020), however, the potential arises for lineage-specific reconstruction of change events, which in turn permits assessment of which parts of a species’ range are “old” *versus* “new” distributional potential.

In this study, we took advantage of these methods, and applied them to a lineage that is admittedly quite simple, particularly as regards temperature niches. That is, the snowfinch species all exist under a single, rather simple, temperature niche (−8–7°C), with the sole exception of *M. nivalis*. This latter species, however, appears to have been able to spread massively westward, to cover much of southern Europe, thanks to one niche expansion event, to a niche of (−9–20°C). Indeed, careful inspection of Fig. 4 suggests that the niche expansion may actually have happened in a subset of the species’ populations: the eastern populations occur dominantly under the ancestral part of the niche, whereas the western populations occur dominantly under the derived part of the niche. In light of this rather recent niche innovation event, one could advance a hypothesis of refugial origin of *M. nivalis*, likely around the Qinghai-Tibet Plateau (Qu et al., 2006; Qu and Lei, 2009; Yang et al., 2006), with subsequent expansion across much of Eurasia, although a detailed phylogeographic study remains to be conducted.

Considerable speculation has revolved around the question of whether niche evolution would elevate or reduce speciation rates (Culumber and Tobler, 2016; Wiens, 2004; Wogan and Richmond, 2015). That is, one could imagine niche evolution allowing a species to overcome distributional barriers, and thereby avoid the population isolation that would lead to speciation. On the other hand, niche evolution might allow the species to “explore” novel distributional situations, which might afford possibilities for colonization of new areas and possible opportunities for speciation. Previous analyses of this question (e.g., (Kozak and Wiens, 2006)) have been compromised by the rampant over-estimation of evolutionary dynamics that inevitably results when availability of areas and associated uncertainty are not considered in ancestral niche reconstructions (Saupe et al., 2018). Using virtual-world simulation approaches, Saupe et al. (2019) found variable effects of niche breadth on speciation rates, with strong effects when climate change is rapid (e.g., abrupt climate change events in glacial-interglacial cycles), but comprehensive analyses have yet to be developed.

In a broader sense, identifying geographic regions and key populations involved in evolutionary changes in ecological niches could prove crucial in identifying target populations for further research aimed at understanding genomic mechanisms underlying niche innovation. That is, functional genomic studies necessarily seek population pairs that are minimally differentiated or separated by independent periods of evolution in isolation (Lamichhaney et al., 2017), so as to avoid a background of evolutionary change in the broader genome. Studies such as this one have considerable potential to make these comparisons more specific—see, e.g., the eastern *versus* western populations of *M. nivalis*, which are closely related, but which may differ in niche characteristics. Another area for future exploration and analysis in other taxa (snowfinch species tend to be broadly sympatric, at least in broad brush-strokes) could be the degree to which areas detected as niche retraction correspond to barriers to gene flow among speciating lineages.

## Acknowledgments

We thank the members of the KUENM working group in the University of Kansas Biodiversity Institute, for their thinking and work over the years.

## Data accessibility

R script and data used in this study, including occurrence data, shapefiles summarizing our hypotheses of accessible areas, and the three phylogenetic trees, are available online at https://doi.org/10.6084/m9.figshare.12915971. The climatic data used in this study can be obtained using the script provided.

## Author Contributions

All authors designed research and compiled data; M.E.C. performed analyses; A.T.P. and M.E.C. wrote the paper; all authors reviewed and approved the manuscript.

